# Validation of Gentamicin and Streptomycin Stability for On-Orbit Microbial Monitoring

**DOI:** 10.1101/2025.05.11.653350

**Authors:** Jordan M. McKaig, Christopher E. Carr

**Author notes:** **Correspondence to:**. Address: ESM Building, Room G10, 620 Cherry St NW, Atlanta, GA 30332, USA.

## Abstract

The Genomic Enumeration of Antibiotic Resistance in Space (GEARS) flight project seeks to characterize the frequency and genomic identity of antibiotic-resistant organisms on the ISS and expand in-space sequencing-based diagnostic capabilities. This project selects for antibiotic-resistant microbes on space station surfaces using agar plates containing antibiotics, which can require storage of these plates for several months. However, little published data is available on the longevity of antibiotics in agar plates. Here, we describe the process of antibiotic selection and validation of antibiotic stability for GEARS. A literature review was conducted on several antibiotics for their intrinsic and acquired resistance in *Enterococcus* and *Staphylococcus* and stability in solution and in agar. Gentamicin and streptomycin were selected for a long-term study tracking changes in bactericidal ability of the antibiotics in agar plates. After 6.5 months of storage at 4°C, we found that both antibiotic types remained stable throughout the test period, although they both exhibited evidence of slight degradation. These findings indicated that either antibiotic would be sufficient for the GEARS study based on stability under cold stowage conditions. Furthermore, this study generated information on long-term stability of antibiotics in agar, which can be useful for microbiology applications beyond spaceflight studies.

## Introduction

Antibiotic resistance has existed for at least tens of thousands of years (Bhullar et al., 2012; D’Costa et al., 2011; Perron et al., 2015; Perry et al., 2016), but it has emerged as a major health risk over the past century. Following the discovery of penicillin in 1928, antibiotics were rapidly and widely harnessed for human and veterinary medicine, livestock, agriculture, and biotechnology. This has revolutionized modern medicine and fundamentally impacted livelihood on Earth, but concurrently has driven extensive proliferation of antibiotic resistance. Today, antimicrobial resistance is identified by the World Health Organization, Centers for Disease Control and Prevention, and the World Medical Association as one of the most prevalent contemporary public health issues (CDC, 2024; WHO, 2019, 2023; WMA, 2022).

Antibiotic-resistant microbes are widespread across external and built environments (Bengtsson-Palme et al., 2017; Larsson & Flach, 2022; Mahnert et al., 2019), and areas that are enclosed and extensively cleaned can actually have a higher prevalence of resistance (Mora et al., 2016). Culture-based methods are commonplace in medicine, agriculture, livestock, and wastewater monitoring, enabling rapid, high-throughput screening for antibiotic-resistant microbes (Caruso, 2018; Haulisah et al., 2021; Kuper et al., 2009; Lemonnier et al., 2023; Manaia et al., 2018; McLain et al., 2016; Reinthaler et al., 2003; Smaill, 2000; Tomida et al., 2013).

Although agar plates containing antibiotics can be used for such methods, little data exists on the long-term longevity of antibiotic stability in agar (over scales of weeks, months, or years). Lyophilized antibiotics are generally stable for months to years, but degrade quickly once reconstituted in water. This is generally not an issue for laboratory settings, as media can be prepared and inoculated with antibiotics as needed. However, certain situations (i.e. remote medicine, spaceflight) require longer-term storage of antibiotics in agar plates if real-time preparation of plates is not feasible, and specific data on the stability of various antibiotics is critical for planning culture-based screening in remote environments.

Over the past few decades of crewed space exploration, the impacts of spaceflight conditions on microbes and potential ramifications on crew health have begun to be realized. In space, microbes can demonstrate increased growth, virulence, pathogenicity, and biofilm formation, as well as changes in metabolism (Huang et al., 2018; Kacena et al., 1999; Kim et al., 2013; Landry et al., 2020; Ott et al., 2020; Rosenzweig et al., 2010; Sharma & Curtis, 2022; Vaishampayan & Grohmann, 2019; Wilson et al., 2007). The unique conditions of spaceflight can likewise exacerbate antibiotic resistance in microbes. This has been characterized in simulated-microgravity experiments on the ground and in culture-based experiments in space (Aunins et al., 2018; Moatti et al., 1986; Tirumalai et al., 2019; Tixador et al., 1985; Urbaniak et al., 2022)

Spaceflight can suppress astronaut immune systems (Sonnenfeld, 1998; Stratis et al., 2023), which increases the risks imposed by these microbial responses to spaceflight. Microbial monitoring is routinely conducted to survey the genomic and functional composition of crewed spacecraft (Yamaguchi et al., 2014). Such monitoring aboard the International Space Station (ISS) has indicated that antibiotic-resistant microbes are present on surfaces proximal to crew members, and may pose risks to astronaut health (Bryan et al., 2021; Urbaniak et al., 2022). Bryan et al. specifically investigated *Enterococcus faecalis* isolates recovered from the ISS, showing that antibiotic-resistant *E. faecalis* exists on-station with the potential for pathogenicity, while also indicating the need to further assess the abundance of antibiotic-resistant microbes on-station.

Genomic Enumeration of Antibiotic Resistance in Space (GEARS) is a NASA- funded spaceflight mission with the goal of characterizing the frequency and genomic identity of antibiotic-resistant microbes on the ISS. This project will use contact slides filled with antibiotic-spiked agar to screen for antibiotic-resistant microbes on internal surfaces on the ISS, and leverage in-space nanopore sequencing instrumentation to identify microbes on-orbit. The GEARS study is one of three complimentary investigations to better understand the adaptation, antibiotic resistance, and adaptation of enterococci in the spaceflight environment. The other two investigations, Enterococcus Growth Advantage Investigation via Tn-seq (EnteroGAIT) and Adaptation and Evolution of Resilient Enterococcus in Space (AERES) are concurrently underway.

To prepare for the GEARS study, it was necessary to first determine which antibiotics would be most useful to generally screen for *Enterococcus*, as the main genus of interest for this project, as well as *Staphylococcus*, another genus with high propensity for antibiotic resistance and human pathogenicity. Next, it was necessary to determine the “shelf life” of these antibiotics once reconstituted in agar, as for GEARS it was important to determine for how long antibiotic stability could be maintained in order to plan mission logistics to prevent significant variability due to loss of antibiotic potency. This manuscript describes the process of antibiotic selection and validation of antibiotic stability and reports results on gentamicin and streptomycin stability over a 6.5 month period.

## Materials and Methods

### Antibiotic selection

A literature review was conducted to identify antibiotics for further study, focusing on two main mechanisms of antibiotic impact related to cell wall biosynthesis (beta-lactams, carbapenems, penicillin-like antibiotics) and ribosome synthesis (aminoglycosides, lincomycin, macrolides, tetracyclines) and associated resistance mechanisms observed in *E. faecalis* (Arias & Murray, 2012). Data on intrinsic and acquired resistance in *Enterococcus* and *Staphylococcus* and long-term (several weeks) stability in solution and in agar plates at 4°C was aggregated from peer-reviewed literature, vendor information, and medical documentation.

Antibiotic recommendations for use in the experimental portion of this study was made based on the presence of intrinsic resistance in *Enterococcus* and *Staphylococcus* and long-term antibiotic viability following reconstitution.

### Contact slide preparation

Three concentrations (high, low, zero) for two antibiotic types (streptomycin, gentamicin) were used in this study. Antibiotic concentrations were determined via preliminary testing on how a model antibiotic-resistant strain *Enterococcus faecalis* OG1RF (ATCC 47077) grew at various antibiotic concentrations. The “high” concentration was the minimum concentration needed to visually attenuate *E. faecalis* OG1RF growth, while the “low” concentration was the minimum concentration needed for *E. faecalis* OG1RF colonies to be visibly smaller than those growing on no-antibiotic TSA. The tested concentrations were 250 µg/mL (high) and 150 µg/mL (low) for gentamicin and 400 µg/mL (high) and 250 µg/mL (low) for streptomycin.

Tryptic soy agar (TSA) (Thermo Scientific R455002) powder was reconstituted in 18 megaohm water following manufacturer instructions, then autoclaved at 121°C for 15 minutes. To ensure that antibiotics were added to agar at a consistent temperature, agar bottles were kept in a water bath at 50°C after autoclaving, for at least 30 minutes and until time of use. Gentamicin (MP Biomedicals 02194530) and streptomycin (MP Biomedicals 0219454125) were reconstituted following manufacturer instructions, then filtered with a 0.2 µm filter. Antibiotic solutions were prepared fresh daily, and stored at 4°C until immediately before use.

Aliquots of antibiotics were added to agar, then agar bottles were swirled gently to mix. Each HYCON contact slide (Millipore 144024) was filled with 12 mL of agar, then was dried in the biological safety cabinet for 30 minutes before closing and transferring to the refrigerator. Contact slides were stored at 4°C in light-shielded conditions for the duration of the stability testing period of 6.5 months.

### Antibiotic stability assay

Approximately once per month, a group of contact slides was removed from storage and their stability was assessed by inoculating with *E. faecalis* OG1RF.

Fresh cultures were prepared and applied to fresh contact slides each month using the following workflow:

A wire loop was flamed and allowed to cool, then was dipped into *E. faecalis* OG1RF stocks stored at-80°C and streaked on three standard TSA plates. Plates were incubated for 24 hours at 37°C. One isolated colony from each plate was used to inoculate a 100 mL flask of tryptic soy broth (TSB). The three flasks were then incubated for 24 hours at 37°C while shaking at 200 rotations per minute (rpm).

Liquid cultures were diluted to 10^-6^, then 50 µL aliquots were plated on contact slides. For each time point, plating was done in triplicate for each group of high/low/zero antibiotic concentration slides (i.e. 3 plates at 250 µg/mL gentamicin, 3 plates at 150 µg/mL gentamicin, 3 plates at 400 µg/mL streptomycin, 3 plates at 250 µg/mL streptomycin, and 6 plates at zero antibiotic). Contact slides were incubated for 48 hours at 37°C.

Following 48 hours of incubation, contact slides were removed from incubation, colony-forming units (CFUs) were counted, and pictures were taken of contact slides from a standardized height above the slides. Images were loaded into ImageJ and the scale was set using the middle line on each contact slide as a reference length, then the ImageJ ROI Manager Tool was used to measure colony diameter.

## Results

### Target antibiotic selection

The literature review focused on 13 antibiotics from 7 classes (FIG.1.), and data on long-term stability in solution and in agar and prevalence of intrinsic and acquired resistance in *Enterococcus* and *Staphylococcus* was collected. These results are shown in TABLE.1. Following this review, gentamicin and streptomycin were selected for further analysis. These antibiotics were among the higher-ranking options for stability in solution, and previous work had indicated that streptomycin is stable in agar for at least 30 days. Furthermore, these specific antibiotics are relevant to the GEARS study – both *Enterococcus* and *Staphylococcus*, the main targets of this project, often display resistance to these antibiotics. Enterococci possess low-level resistance to aminoglycosides overall (a category that includes both gentamicin and streptomycin), and can further acquire high-level resistance. Furthermore, staphylococci can develop resistance to both antibiotics.

**FIG. 1.**
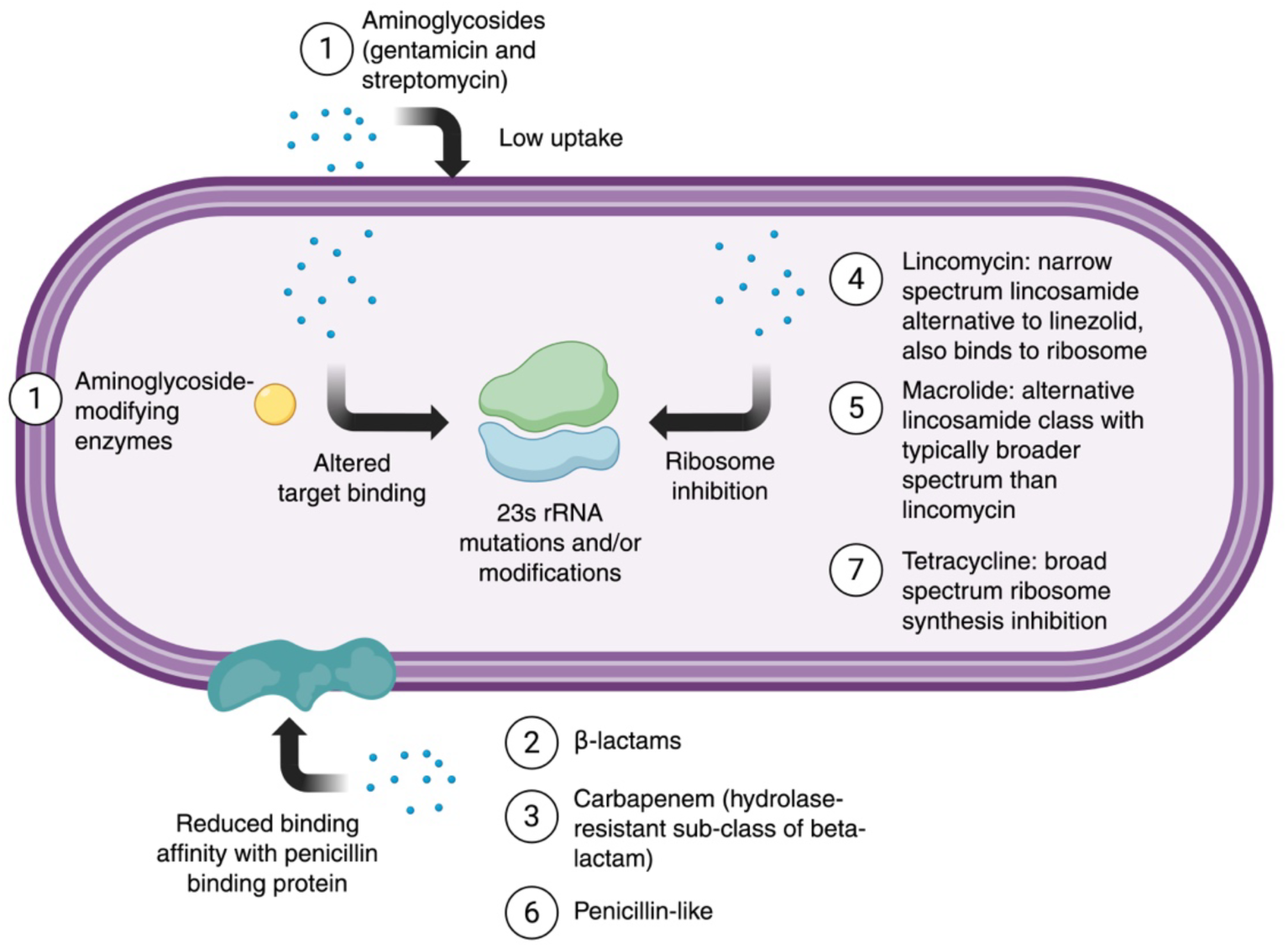
Mechanisms of resistance for antibiotics included in literature review. Antibiotics are numbered based on order in TABLE.1. Adapted from Arias & Murray 2012.

**TABLE. 1.** Antibiotic literature review results. Antibiotics are organized alphabetically within their respective classes. Stability in solution and in agar at 4°C is given, then resistance in *Enterococcus* and *Staphylococcus* is briefly described.

| Class | Antibiotic | Stability in solution at 4°C | Stability in agar plates at 4°C | Enterococcus resistance |  | Staphylococcus resistance |
| --- | --- | --- | --- | --- | --- | --- |
|  |  |  |  | Low-level | High-level |  |
| Aminoglycoside | Amikacin | 60 days<br>(World Health Organization, 2014) | Unknown | Intrinsic<br>(Chow, 2000) | Can be acquired<br>(Chow, 2000) | Can be acquired<br>(Gold et al., 2014) |
|  | Gentamicin | 4 weeks<br>(Bastani et al., 2005) | Unknown | Intrinsic<br>(Hollenbeck & Rice, 2012) | Can be acquired<br>(Hollenbeck & Rice, 2012) | Intrinsic<br>(Cattoir, 2016; Domínguez et al., 2002; Vestergaard et al., 2016) |
|  | Kanamycin | 12 months<br>(US Biological Life Sciences, 2023) | 30+ days<br>(Ryan et al., 1970) | Intrinsic<br>(Hollenbeck & Rice, 2012) | Unknown | Observed, unknown if intrinsic or acquired<br>(Domínguez et al., 2002) |
|  | Streptomycin | 2 weeks<br>(Millipore<br>Sigma, 2021) | 30+ days<br>(Ryan et<br>al., 1970) | Intrinsic<br>(Hollenbeck<br>& Rice,<br>2012) | Can be<br>acquired<br>(Chow,<br>2000) | Intrinsic (Jensen<br>& and Lyon,<br>2009) |
|  | Tobramycin | 96 hours<br>(GlobalRPH,<br>2001) | Unknown | Intrinsic<br>(Hollenbeck<br>& Rice,<br>2012) | Unknown | Observed,<br>unknown if<br>intrinsic or<br>acquired<br>(Domínguez et<br>al., 2002) |
| Beta-lactam | Cephalosporin | 7-30 days<br>(Xu et al.,<br>2002) | Unknown | Intrinsic<br>(Hollenbeck<br>& Rice,<br>2012) | Intrinsic<br>(Hollenbeck<br>& Rice,<br>2012) | Intrinsic (Bruns<br>& Keppeler,<br>1980) |
| Carbapenem<br>(beta-lactam with<br>hydrolase<br>resistance) | Ertapenem | 24 hours<br>(GlobalRPH,<br>2008) | Unknown | Intrinsic<br>(Werth,<br>2022) | Unknown | Can be acquired<br>(Gesser et al.,<br>2004; Ghosh &<br>Banerjee, 2016) |
| Lincomycin | Clindamycin | 32 days<br>(GlobalRPH,<br>2007) | Unknown | Intrinsic<br>(Hollenbeck<br>& Rice,<br>2012) | Unknown | Can be acquired<br>(Bastani et al.,<br>2005) |
| Macrolide | Erythromycin | 7 days (Al-Ramahi et<br>al., 2015) | 30+ days<br>(Ryan et<br>al., 1970) | Can be<br>acquired<br>(Kristich et<br>al., 2014) | Can be<br>acquired<br>(Kristich et<br>al., 2014) | Can be acquired<br>(Mikłasińska-Majdanik, 2021) |
| Penicillin-like<br>beta-lactams | Amoxicillin | 10 days (Al-Ramahi et<br>al., 2015) | Unknown | Can be<br>acquired<br>(Conceição<br>et al., 2012) | Can be<br>acquired<br>(Conceição<br>et al., 2012) | Intrinsic (Bruns<br>& Keppeler,<br>1980) |
|  | Ampicillin | 3 weeks<br>(Sigma-Aldrich, 2016) | 7 days<br>(Ryan et al., 1970) | Can be acquired<br>(Hollenbeck & Rice, 2012; Kristich et al., 2014) | Can be acquired<br>(Hollenbeck & Rice, 2012; Kristich et al., 2014) | Intrinsic (Bruns & Keppeler, 1980) |
|  | Penicillin | 2 weeks (21) | 7 days<br>(Ryan et al., 1970) | Intrinsic<br>(Hollenbeck & Rice, 2012) | Unknown | Intrinsic<br>(Bruns & Keppeler, 1980; Lowy, 2003) |
| Tetracycline | Tetracycline | 2 weeks<br>(Barrick Lab, 2022) | 30+ days<br>(Ryan et al., 1970) | Can be acquired<br>(Wilcks et al., 2005) | Can be acquired<br>(Wilcks et al., 2005) | Can be acquired<br>(Grossman, 2016) |

### Antibiotic stability over 6.5 month test period

During each testing interval, colony-forming units (CFUs) were counted on each set of contact slides (high/low gentamicin and streptomycin, no antibiotic). FIG.2. displays the CFU counts over 6.5 months. During the testing period, no CFUs ever grew on the high-concentration contact slides for both antibiotic types.

**FIG. 2.**
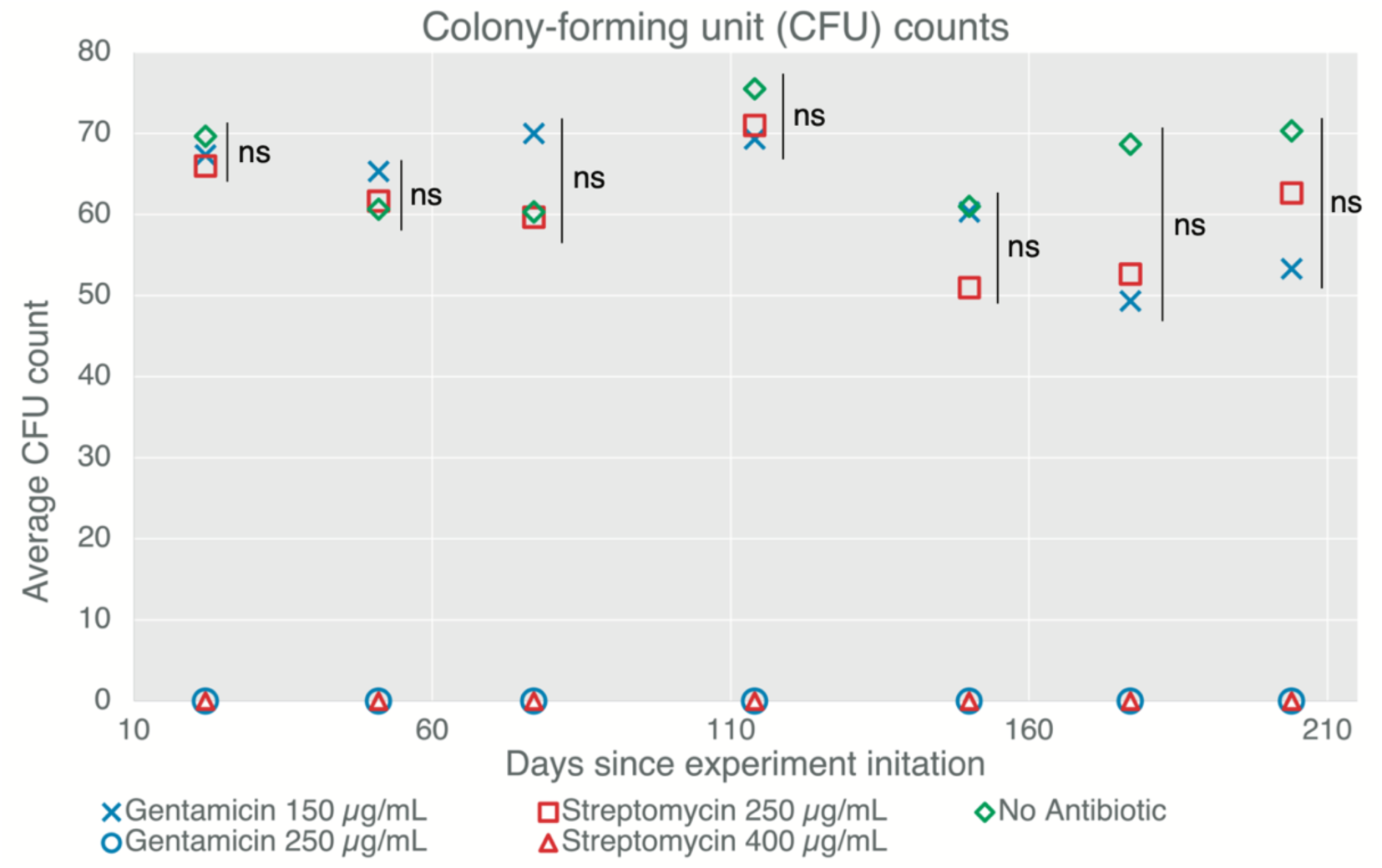
Average colony-forming unit (CFU) counts for contact slides containing no antibiotic, low-concentration and high-concentration gentamicin, and low-concentration and high-concentration streptomycin over 6.5 months. For each time point, one-tailed t-tests were performed on the average CFU counts for the contact slides containing no antibiotic and low-concentration gentamicin and streptomycin. The asterisks denote level of statistical significancy (no significance: ns; p < 0.05: *; p < 0.01: **; p < 0.001: ***; p < 0.0001: ****).

Additionally, no statistically significant difference was found in the colony counts between the no-antibiotic, low-gentamicin, and low-streptomycin plates for each time period.

Colony diameter was measured as a general indicator of antibiotic stability. This was intended to serve as an additional exploration of a potentially useful but currently unvalidated method to further inform antibiotic selection (see Discussion for further details). To account for variations in number of colonies and incubation conditions, the colony diameters for each contact slide were averaged, then the average diameters for each of type of contact slide were averaged. Finally, the average colony diameter for each antibiotic type were compared to the average colony diameter for the no-antibiotic contact slides for each time point. FIG.3. displays these results over the 6.5 month test period.

**FIG. 3.**
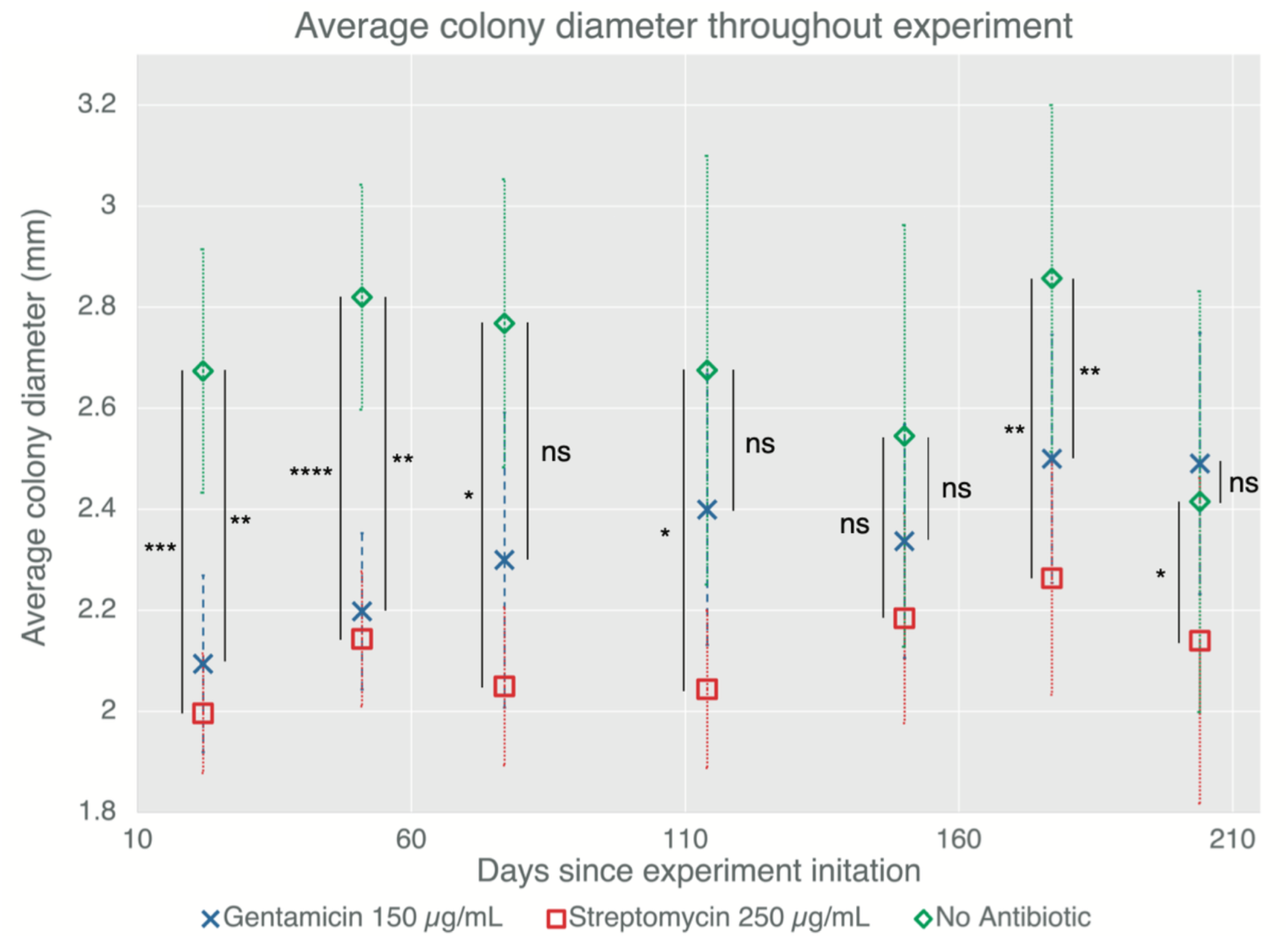
Average colony diameter on contact slides containing no antibiotic, low-concentration gentamicin, and low-concentration streptomycin over 6.5 months. To account for variations in number of colonies and incubation conditions, the colony diameters for each contact slide were averaged, then the average diameters for each of type of contact slide were averaged. Finally, the average colony diameter for each antibiotic type were compared to the average colony diameter for the no-antibiotic contact slides for each time point. For each time point, one-tailed t-tests were performed between the average colony diameter for the no-antibiotic contact slide (pink) and the average colony diameters for the low-concentration gentamicin (green) and streptomycin (blue) contact slides. The asterisks denote level of statistical significancy (no significance: ns; p < 0.05: *; p < 0.01: **; p < 0.001: ***; p < 0.0001: ****).

Throughout the duration of testing, the average colony diameter for antibiotic contact slides was lower than that of the no-antibiotic contact slides, with varying levels of statistical significance (or lack thereof). The only exception to this is the last time point, where the average diameter on the gentamicin contact slides was larger than the no-antibiotic contact slides, although the difference was not statistically significant. This could be an indicator of slight gentamicin degradation, although longer-duration experimentation would have been necessary to further resolve this trend.

## Discussion

In this study, we present data on the long-term stability of gentamicin and streptomycin in agar, which was used to inform antibiotic selection and planning for the GEARS spaceflight mission. We discuss the criteria for selecting test antibiotics, show that both tested antibiotics maintain stability over the 6.5 month test period, and discuss a novel method for using colony diameter to gain additional insight into antibiotic stability.

Through the literature review, gentamicin and streptomycin were selected for further validation for the GEARS study, due to their viability in solution and agar plates and prevalence of resistance to these drugs in *Enterococcus* and *Staphylococcus*. *Enterococcus* possess low-level intrinsic resistance to aminoglycosides (a class that includes both gentamicin and streptomycin), based on poor drug uptake and drug inactivation by *Enterococcus-*produced enzymes (Miller et al., 2014). *Staphylococcus* similarly exhibit low-level intrinsic resistance to gentamicin (Cattoir, 2016; Vestergaard et al., 2016) and exhibit widespread emergent low-level resistance to streptomycin (Jensen & and Lyon, 2009). Additionally, streptomycin has been shown to be viable in agar plates for at least 30 days (Ryan et al., 1970), and gentamicin and streptomycin are viable in solution for 4 weeks (Bastani et al., 2005) and 2 weeks (Millipore Sigma, 2021) respectively. To provide flexibility for mission logistics, we evaluated whether antibiotics could remain stable in agar plates in cold stowage conditions for up to 6.5 months. The “shelf life” would limit the time between preparation and usage, and since GEARS plates are being returned to Earth, would imply limits on the time between launch and landing to avoid overgrowth of plates. These two antibiotics were subjected to additional stability testing in order to inform GEARS flight planning.

Perhaps the most prominent general finding from the literature review was the deep lack of data on the stability of antibiotics in agar. This is not surprising: conventional laboratory operations allow for rapid preparation of antibiotics for use in agar or solution, and general best practices for antibiotic resistance research dictate using freshly-prepared antibiotics. However, it is useful to have information on long-term stability of antibiotics in agar for applications where the tools of a standard microbiology lab may not be accessible (e.g. spaceflight, remote medicine, livestock monitoring). By validating the long-term stability of our antibiotics of interest for the GEARS study, we collected data that may be useful for other situations where antibiotic resistance monitoring needs to be conducted but a constant production of antibiotic-containing consumables is not feasible. In addition, these findings may have implication for the natural use of antibiotics by microbes; extended stability of antibiotics could support ecological functions such as niche maintenance, defense, or shape inter-species interactions.

Throughout the 6.5 month testing period, there were no significant differences between CFU counts on low-antibiotic and no-antibiotic contact slides, although the colonies on the low-antibiotic contact slides were visibly smaller than those of the no-antibiotic contact slides. Additionally, no colonies ever grew on the high-antibiotic contact slides. This indicates that the chosen test concentrations were well-suited for growing the test model organism on antibiotic concentrations that partially and fully attenuated growth, and that antibiotics remained stable enough to have these effects throughout the test period.

Antibiotic decay was estimated to be no greater than 40% for gentamicin and 37.5% for streptomycin as described below. No colonies were observed on the high-concentration antibiotic contact slides (250 µg/mL for gentamicin and 400 µg/mL for streptomycin) throughout 6.5 months of storage, while colonies were observed on the low-concentration antibiotic contact slides (150 µg/mL for gentamicin and 250 µg/mL for streptomycin) at the initiation of the experiment, when the contact slides were freshly made. This implies that the high-concentration antibiotic contact slides did not decay down to an equivalent functional concentration of the low-concentration antibiotic contact slides during the testing period. Thus, the remaining functional antibiotic doses must be greater than 150/250 or 60% of the original for gentamicin, and 250/400 or 62.5% for streptomycin.

Based on these findings, it was determined that either antibiotic option would be suitable for the GEARS study when targeting 30-45 day duration Commercial Resupply Services (CRS) missions. Ultimately, gentamicin was selected, due to satisfactory stability and anecdotal observations of streptomycin interfering with downstream DNA extraction and sequencing, although at tested doses higher than considered for culturing.

In this experiment, we used colony diameter as an additional proxy metric for antibiotic efficacy. This was based off an observation in preliminary testing that colony diameter decreased with increasing antibiotic concentration. Thus, we decided to record colony diameter to explore how it changed as antibiotics decayed over the test period. It is important to caution that this is not a validated method for directly measuring antibiotic efficacy, and is intended to be a potentially-useful metric that enriches our conclusions and may merit further exploration. Further testing would be necessary to determine whether there is a quantifiable relationship between antibiotic concentration/efficacy and colony diameter, and to explore how this relationship may translate into clinical applications. These results imply that for longer duration CRS missions, up to six months, some degradation may be present, but does not exceed 40%, and could be lower.

The GEARS study is currently underway, with the first launch and operations having occurred in March and April of 2024, and the second in late 2024. It is possible that this mission will reveal additional unexpected information about the utility of antibiotics under the unique conditions of spaceflight (i.e. launch conditions, radiation, and microgravity). As space missions increase in duration, distance from Earth, and size of crew, it will be critical to continue to update understanding of the microbes that coexist with the crew, which can have drastic implications on astronaut health.

## Acknowledgements

The authors would like to thank the GEARS team including Project Scientist Gwo-Shing Sun, Co-PI Sarah L. Castro-Wallace, Sarah Stahl-Rommel, Christian Mena, Hang N. Nguyen, Christian Castro, G. Marie Sharp, Noelle C. Bryan, Ralf Moeller, Elisabeth Grohmann, and Co-PI Aaron S. Burton. Figure 1 was created using BioRender.com. This material is based upon work supported by the National Aeronautics and Space Administration under Grant No. 80NSSC21K0234 to C.E.C.

